# Human Mesofluidic Intestinal Model for Studying Transport of Drug Carriers and Bacteria Through a Live Mucosal Barrier

**DOI:** 10.1101/2024.09.18.613692

**Authors:** Chia-Ming Wang, Hardeep S. Oberoi, Devalina Law, Yuan Li, Timothy Kassis, Linda G. Griffith, David T. Breault, Rebecca L. Carrier

## Abstract

The intestinal mucosal barrier forms a critical interface between lumen contents such as bacteria, drugs, and drug carriers and the underlying tissue. Current *in vitro* intestinal models, while recapitulating certain aspects of this barrier, generally present challenges with respect to imaging transport across mucus and uptake into enterocytes. A human mesofluidic small intestinal chip was designed to enable facile visualization of a mucosal interface created by growing primary human intestinal cells on a vertical hydrogel wall separating channels representing the intestinal lumen and circulatory flow. Type I collagen, fortified via cross-linking to prevent deformation and leaking during culture, was identified as a suitable gel wall material for supporting primary organoid-derived human duodenal epithelial cell attachment and monolayer formation. Addition of DAPT and PGE2 to culture medium paired with air-liquid interface culture increased the thickness of the mucus layer on epithelium grown within the device for 5 days from approximately 5 mm to 50 μm, making the model suitable for revealing intriguing features of interactions between luminal contents and the mucus barrier using live cell imaging. Time-lapse imaging of nanoparticle diffusion within mucus revealed a zone adjacent to the epithelium largely devoid of nanoparticles up to 4.5 hr after introduction to the lumen channel, as well as pockets of dimly lectin-stained mucus within which particles freely diffused, and apparent clumping of particles by mucus components. Multiple particle tracking conducted on the intact mucus layer in the chip revealed significant size-dependent differences in measured diffusion coefficients. E. coli introduced to the lumen channel were freely mobile within the mucus layer and appeared to intermittently contact the epithelial surface over 30 minute periods of culture. Mucus shedding into the lumen and turnover of mucus components within cells were visualized. Taken together, this system represents a powerful tool for visualization of interactions between luminal contents and an intact live mucosal barrier.

## Introduction

The gastrointestinal tract is lined with mucus, a consistently renewed barrier protecting the underlying tissue against pathogen invasion and controlling the diffusion of luminal contents to the intestinal epithelium(1-3). As oral drug delivery remains the most commonly used and preferred route of drug administration, and intestinal mucus is considered a potential major biological barrier to drug and drug carrier permeation and uptake, studying the interactions between mucus, drugs, and drug carriers can provide insight into important design features of drug delivery systems. Further, understanding interactions between the gut microbiota and mucus is crucial for unraveling the complex host-microbe relationship. In recent years, the gut microbiota has been extensively linked to a broad spectrum of human diseases and disorders, such as obesity(4), inflammatory bowel diseases(5), diabetes(6), and neurological disorders(7). Mucus-bacteria interactions are increasingly recognized to be critical to the impact of the microbiota on human health and diseases(8). Changes in the mucus barrier have been observed in disease states associated with gut inflammation and infection(9), suggesting an altered mucus barrier might contribute to disrupted gut-microbiota homeostasis through currently poorly understood mechanisms.

Animal models(10), excised tissues(11, 12), and collected native mucus(3, 13-16) are commonly used to study interactions of lumen contents with intestinal mucus(17). These systems provide the advantage of preserving the complexity of mucus composition, including cellular components (e.g., DNA)(16) significant to mucus barrier properties, and offer the ability to access mucus from various segments of the gastrointestinal tract or from different disease models to study associated variations in mucus properties. However, interspecies differences in mucosal biology and non-isotropic properties of intestinal mucus motivate the development of a human *in vitro* intestinal model, enabling facile investigation of an intact mucus layer on intestinal tissue. Conventional 2-dimensional (2D) *in vitro* intestinal models commonly involve culturing human cell lines (e.g., Caco-2 cells) or, more recently, organoid-derived human primary intestinal cells on a porous cell culture insert membrane and have been extensively utilized to study the intestinal mucosal barrier(18-21). These 2D *in vitro* intestinal models can incorporate various components such as immune cells(22) and commensal bacteria(23) to mimic the human intestine better and provide the ability to study the interaction of luminal contents with mucosal barrier in a controlled environment. Over the past decade, advances in microfabrication and microfluidic techniques have facilitated the emergence of gut-on-a-chip models(24-26). These innovative systems offer a more physiologically relevant platform to study the interactions between the mucosal barrier and luminal contents. Unlike conventional models, gut-on-a-chip models can replicate key aspects of the gut environment, including lumen and circulatory flow(27-29), enabling longer term investigation of host-microbe interactions(30, 31). Most gut-on-a-chip systems, as well as conventional cell culture insert-based models, involve culturing cells on a horizontal membrane, limiting the ability for visualization of the mucosal-lumen interface and the interaction of lumen contents with the mucus barrier using standard microscopy without sacrificing the culture and complicated sample processing. Further, the mucus layer produced on most intestinal models is markedly thinner than that which exists *in vivo*. Caco-2 cells do not secrete mucus but are frequently cultured with goblet cell-like HT29-MTX cells, and these co-cultures produce a mucus layer approximately 5 um thick(32, 33). Primary ileal and rectal intestinal cultures on the Transwell system have been reported to produce mucus layers approximately 25 and 35 um thick, respectively(18), which is still markedly lower than the mucus layer thickness *in vivo* (200-700 μm depending on the intestinal sections)(34).

In this study, we report a new mesofluidic gut-on-a-chip model with organoid-derived primary human intestinal cells grown on a vertical gel wall, enabling facile visualization of the mucosal interface using a standard confocal microscope. The media composition and culture techniques were varied to enable the production of a mucus layer of thickness comparable to that of the intestine *in vivo*. The ability to conduct real-time analysis of the transport of nanoparticle drug carriers and bacteria through the mucus and of proximity to the epithelial apical surface in live cultures was demonstrated. Further, the turnover of the mucus layer including cellular internalization of mucus components was directly visualized. We demonstrate that this mesofluidic gut model is a powerful tool to capture real-time interactions between mucus and luminal contents (i.e., bacteria and drug carriers) in a defined, well-controlled environment. This tool can be applied for *ex-vivo* screening of formulations and fundamental studies of how drug carrier and bacteria properties influence their permeation through mucus.

## Experimental

### Mesofluidic gut chip design and fabrication

Mesofluidic gut chips were fabricated from PDMS polymer (polydimethylsiloxane; Sylgard™ 184, Krayden DC4019862) using a soft-lithography technique. The lumen and side channels were designed with similar dimensions (both 500 μm high x 2 mm wide) but with different volumes due to the difference in length and channel shape: lumen channel was 70 μl and side channel was 40 μl. A gel wall separating the lumen and side channels was formed via introduction of gel precursor material as described below and was flanked by a ridge (300 μm high starting from the top) on each side extending from the top that acted as phase guides supporting the hydrogel wall. To fabricate the PDMS portion of the chip, PDMS prepolymer (12:1 w/w ratio of PDMS to curing agent) was cast on a 3-D printed mold of the inverse channel design and degassed in a desiccator chamber for 1 hour, followed by curing at 60°C for 5 hours. After peeling the PDMS device off the mold, the inlet, outlet, and gel loading holes were punched using EMS-Core sampling tools (1.2 mm diameter for the inlet and outlet and 0.5 mm diameter for the gel loading holes; Electron Microscopy Sciences 69039-12 and 69039-05). The surfaces of the PDMS device and a glass coverslip (Superslip^®^ Micro Glasses, 24 × 50 mm, No.1, Electron Microscopy Sciences 72236-50) were subjected to plasma treatment for 40 seconds using Harrick Plasma Cleaner (PDC-001), and the plasma-treated surfaces of the PDMS device and glass coverslip were placed in contact for binding. The mesofluidic gut chip was pressed under a vise while heated at 75°C for 2 hours and then overnight (12-16 hours) at room temperature to enhance adherence prior to introduction of gel precursor to form the gel wall cell culture surface.

### Preparation of cross-linked collagen gel wall

A 2 mg/ml neutralized type I collagen (pH=7.2-7.4) solution was prepared by diluting 5 mg/ml of rat tail type I collagen (Cultrex 3-D Culture Matrix, R&D Systems™ 344702001) with neutralization buffer(20). The neutralization buffer is composed of HEPES (20 mM, Gibco™ 15630080), NaOH (9.4 mM), and NaHCO_3_ (53 mM, Sigma-Aldrich S5761) in phosphate buffered saline (PBS, Fisher BioReagents BP661-10). The neutralized collagen mixture was centrifuged at 2500 rpm for 5 minutes at 4°C, and 21 μl of the supernatant was added into the stromal “gel guide channel”, followed by incubation at 37°C for 1 hour for gelation. The collagen gel wall formed due to surface tension effects as the gel precursor interacted with the phase guide ridges. PBS was added into both lumen and side channels, and the chip was placed at 4°C for 30 minutes. A gradient cross-linked collagen gel wall was prepared using modification of a previously published protocol for cross-linking collagen substrates on Transwell inserts(35, 36). Briefly, PBS in the side channel was replaced with MES buffer (0.1M, pH=5.5; Sigma-Aldrich 76039) which contained 353 mM 1-Ethyl-3-(3-dimethylaminopropyl) carbodiimide (EDC; Thermo Scientific 22980) and 88 mM N-Hydroxysuccinimide (NHS; Sigma-Aldrich 130672) to initiate cross-linking of the collagen gel wall from the side channel moving inward toward the lumen. The chip was placed at 4°C for 40 minutes during cross-linking, and cross-linking reagents as well as the PBS in the side channel were replaced with DI water. The chip was stored at 4°C prior to sterilization. To sterilize the gut chip, 70% ethanol was introduced to both lumen and side channels, and the whole device was immersed in 70% ethanol for 30 seconds. The whole device was then transferred to a petri dish filled with sterile water to wash off ethanol from the surface. The lumen and side channels were rinsed three times using PBS and filled with PBS. The chip was placed in a humidifying box (a tip box filled with sterile water in the bottom compartment) at 37°C to avoid the collagen gel wall drying up prior to cell seeding.

### Human duodenal organoid culture and human mesofluidic duodenal chip establishment

Primary human duodenal epithelial cells used in this study were derived from duodenal organoids (donor H416) obtained from the Harvard Digestive Diseases Center (HDDC) Organoid Core (David Breault Lab) at Boston Children’s Hospital. Duodenal crypt tissues were obtained by endoscopic tissue biopsies from grossly normal appearing regions of the duodenum in patients upon the donors’ informed consent to establish duodenal organoid cultures. Duodenal organoids were expanded in Matrigel^®^ matrix (Corning™ 354234) with duodenal expansion media (EM, Table 1) and passaged every 7 days. To establish human mesofluidic duodenal chip cultures, 4% Matrigel^®^ and 50 μg/ml of rat tail collagen type I (Corning™ 354236) in Dulbecco’s PBS supplemented with CaCl_2_ and MgCl_2_ (DPBS; Gibco™ 14040117) were introduced into the lumen channel and incubated for 2 hours at 37°C, followed by rinsing with EM once prior to cell seeding. Duodenal organoids (passage 15-20) were collected on day 7 of culture with media together by physically scraping organoid containing Matrigel^®^ matrix (25 μl of Matrigel^®^ for each droplet) off the tissue culture plate, followed by centrifugation (500g x 5 min, 4°C) of the collected Matrigel^®^ and culture medium to obtain the cell pellet. The resulting organoid pellet was resuspended with 3 ml of 0.5 mM disodium ethylenediaminetetraacetic acid (EDTA; Sigma-Aldrich E6635) in PBS to dissolve Matrigel^®^ and pelleted again by centrifugation (500g x 5 min, 4°C). The organoid pellet was resuspended and trypsinized using 1 ml of 0.25% trypsin-EDTA (Gibco™ 25200072) for 5 min in 37°C water bath and dissociated into single cells by manually pipetting with a bent tip. Trypsin was neutralized with 5 ml of Advanced Dulbecco’s Modified Eagle Medium (DMEM; Gibco™ 12491015) supplemented with 10% fetal bovine serum (FBS, R&D Systems S11150), 1x GlutaMAX (Gibco™ 35050061), and 1% Penicillin-Streptomycin (Gibco™ 15140122), followed by centrifugation (300g x 5 min, 4°C) to obtain the cell pellet. Finally, duodenal stem cells were resuspended in EM for seeding. Primary human duodenal stem cells were manually pipetted into the lumen channel at a density of 1.43 × 10^6^cells per ml (or 1.42 × 10^6^cells per cm2 with respect to the surface area of the collagen gel wall), and the duodenal chips were cultured tilted at 90° angle such that cells settled onto the surface formed by the gel wall lining the lumen channel throughout the culture period. The duodenal chips were fed by replacing lumen and side channel contents with fresh EM every day by manually pipetting for 3-5 days until confluent monolayer formation, followed by differentiation using differentiation media (DM) for 5 days before use in experiments. Differentiation medium formulation optimization and air-liquid interface differentiation studies were conducted using the same cell culture procedure described above but used varying differentiation medium formulations and differentiation methods. Cell culture medium formulations used are detailed in Table 1.

**Table 1.**
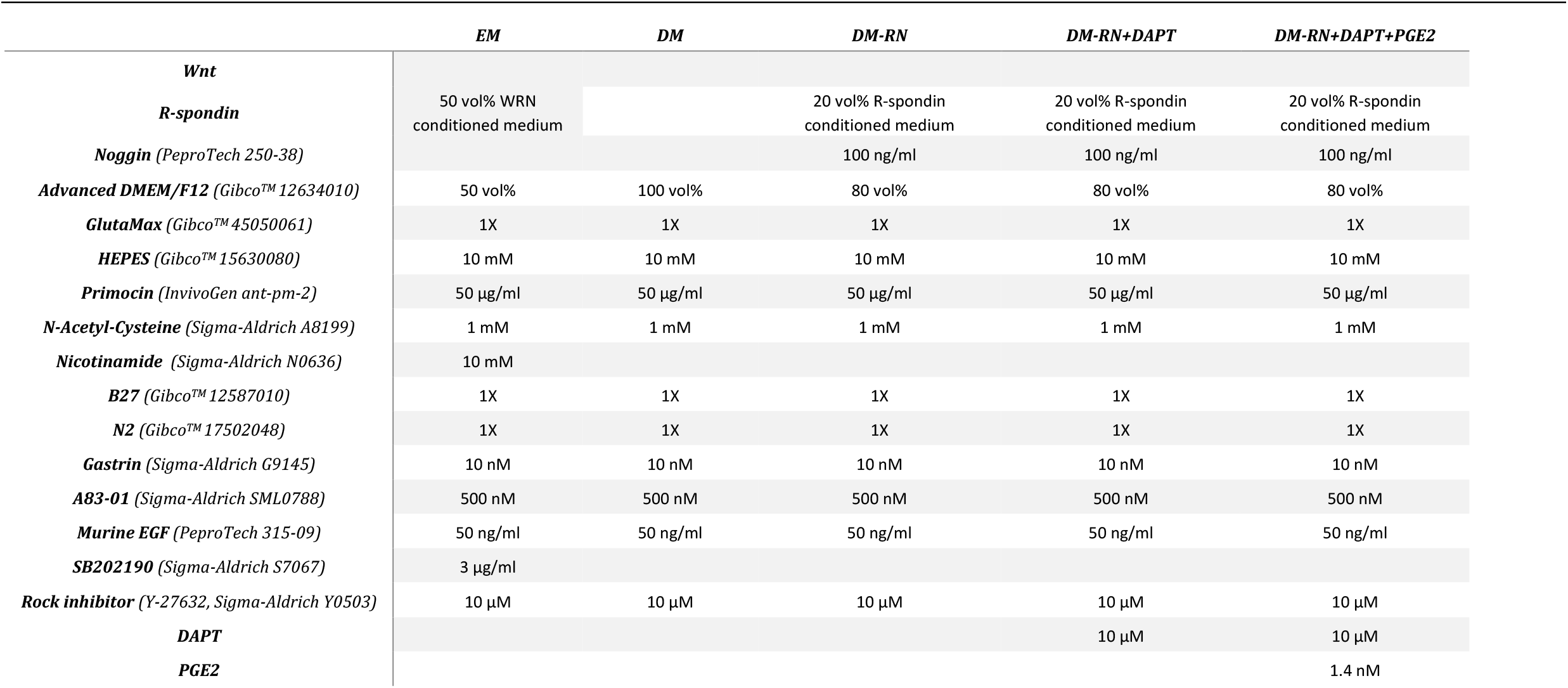
Cell Culture Media Formulations.

### Live cell staining

Live cell staining with fluorescently conjugated Wheat Germ Agglutinin (WGA; Invitrogen™ W32466) for labeling sialic acids and N-acetylglucosamine in mucus and Hoechst 33342 (Thermo Scientific™ 62249) for labeling cell nuclei was conducted according to previously published protocols(37). A solution of WGA (10 μg/ml), Hoechst 33342 (5 μg/ml), and 200-nm carboxylate-modified fluorescent polystyrene microspheres (diluted to 0.0025% solids, Invitrogen™ F8811,) was prepared in corresponding differentiation media and added into the lumen channel, followed by incubation at 37°C for 30 minutes. A ZEISS LSM800 confocal microscope was utilized to collect fluorescence images of the mucosal interface with the same imaging parameters used for each sample from the same set of experiments.

### Alcian Blue mucin quantification assay

Alcian blue was used as a colorimetric assay to quantify mucin concentration in medium samples(38). Medium samples consisted of fluid collected from the lumen channel of human duodenal chips at the end of the cell culture period before imaging. They were stored at -20°C prior to use. Serial dilutions of mucin type II from porcine stomach (Sigma-Aldrich M2378) in DI water with concentrations ranging from 0 to 500 μg/ml were used to generate a standard curve. Medium samples and standards were added to a 96-well plate, mixed with Alcian Blue solution (1% in 3% acetic acid, pH 2.5; Sigma-Aldrich B8438) at a 3:1 ratio and equilibrated at room temperature for 2 hours, followed by centrifugation at 1870 g for 30 minutes to separate the precipitates. The precipitates were then washed once using resuspension buffer, and the resulting suspension was centrifuged at 1870g for 30 minutes to obtain mucin pellets. The resuspension buffer is composed of 40% v/v ethanol and 25 mM magnesium chloride (Sigma-Aldrich M8266) in 0.1M sodium acetate buffer (Fluka™ 15633190). The mucin pellets were resuspended in 10% w/v sodium dodecyl sulfate (SDS; Sigma-Aldrich 436143) solution and subjected to absorbance measurement at 620 nm using a microplate reader (BioTek^®^ PowerWave XS). Mucin concentrations were calculated using the absorbance values and the standard curve absorbance-mucin concentration linear relationship.

### Preparation of drug-rich amorphous nanoparticles (ANPs)

A controlled solvent-antisolvent precipitation method was used to prepare a dilute suspension of amorphous nanoparticles containing an active pharmaceutical ingredient (API), as described previously(39). Briefly, a solvent phase (acetone) containing the API “ABT-450” (AbbVie Inc.), polymer and surfactant was mixed with an aqueous phase (anti-solvent, DI water) at a controlled rate using an impinging jet (IJ) mixer, resulting in the formation of a colloidal suspension. The IJ mixer was Y-type with a 10° offset and having diameters of 0.6 mm for the solvent and 2.75 mm for the anti-solvent arms. The organic phase consisted of 80% w/w acetone (Sigma-Aldrich), 10% w/w DI water, 7% w/w ABT-450, 2% w/w Copovidone (Ashland), 1% w/w SDS (Sigma-Aldrich), and 0.1% w/w fluorescein-5-isothiocyanate (FITC; Invitrogen™ F143). The liquids were pumped using NE-4000 programmable syringe pumps (New Era Pump Systems Inc.) into the IJ mixer at varying organic to aqueous phase flow rate ratios: 1. 1 to 6 (8 ml/min to 48 ml/min) flow rate ratio creating small ANPs (125 nm) and 2. 1 to 4 (8 ml/min to 32 ml/min) flow rate ratio creating large ANPs (230 nm). The resulting nanosuspension was subjected to acetone removal using a rotary evaporator with the water bath temperature at 30°C. Finally, particle size and zeta potential measurements were performed on the resulting nanosuspension using Anton Paar Litesizer 500. The nanosuspension was diluted with PBS to 0.0025% w/w colloids prior to analysis. Particle size and zeta potential analyses were performed at room temperature and at least three separate measurements were conducted.

### Native porcine intestinal mucus collection and mucus sample preparation

Native porcine intestinal mucus was collected from porcine jejunum shortly after animal sacrifice at a local abattoir (Research 87 Inc., Boylston, MA) and delivered within 2 hours. The pig jejunum was gently rinsed with cold water to remove bulk waste before mucus collection. The mucus layer was gently scraped from the tissue using a metal spatula and stored in microcentrifuge tubes at -80°C prior to use. Frozen pig intestinal mucus was thawed at room temperature for 30 minutes before particle tracking experiments were conducted. Thawed pig intestinal mucus (150 μl) was added to an 8-well Nunc™ Lab-TeK™ Chamber Slide system (Thermo Scientific™ 177410). Next, diluted ABT-450 ANPs (7.5 μl, 0.0025% w/w colloids in PBS) were vortexed, sonicated to mix, and then gently pipetted dropwise on the top of mucus samples with minimum disturbance to the mucus surface. Samples were incubated in the dark in a humidifying chamber at room temperature for 2 hours to allow particle diffusion without additional mixing before video collection for particle tracking.

### Multiple particle tracking (MPT)

Particle movement videos were recorded using an inverted Olympus IX51 microscope with an attached Olympus DP70 digital color camera and an X-Cite 120 fluorescence illumination system. 20-second videos were collected at a frame rate of 29 fps. Tracking videos were taken within the mucus layer where mucus was labeled and visualized by WGA.

### Particle diffusion analysis

Tracking videos were analyzed frame by frame using a MATLAB algorithm to obtain x- and y-coordinate trajectories of diffusing particles over time(40). With the trajectory data, mean squared displacement (MSD) and effective diffusivity can be calculated using the following equations:

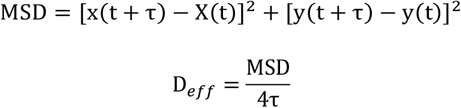

where, x(t) and y(t) represent the particle x- and y-coordinate position at a given time point and τ is the time scale. D_eff_ is used to quantitatively characterize the particle diffusion through mucus at a given time scale. Calculated D_eff_ can be used to estimate the fraction of a population of particles expected to penetrate through a mucus layer of given thickness over time using numerical integration of Fick’s second law:

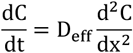

where C is the normalized particle concentration defined as C_p_ (particle concentration)/C_p0_ (bulk particle concentration in the lumen channel), t is the time, and x is the x-coordinate position. The following initial and boundary conditions of C(x,t) are used for prediction: The constant particle concentration in the lumen channel was defined as C(0,t)=1, for t>0 as boundary conditions, and the initial particle concentration present at the mucus layer front was defined as C(x,0)=1 when x=0, otherwise, C(x,0)=0. The fraction of particles predicted to penetrate through the intact mucus layer in the mesofluidic duodenal chip was calculated from a solution to Fick’s second law subject to initial and boundary conditions provided above(3), with the mucus layer thickness defined as 50 μm and the time scale defined as 1.5 hour.

### Bacteria preparation and co-culture with human duodenal chips

*Escherichia coli* (*E. coli* MG1655) and *Lactobacillus rhamnosus* (*L. rhamnosus*, KLE2101, gift from Kim Lewis lab at Northeastern University) were used as model bacteria to study interactions of bacteria with the mucus barrier in the mesofluidic gut model. *E. coli* was transformed with a green fluorescent protein (GFP) plasmid (p-mut2-GRP, gift from Marcia Goldberg at Massachusetts General Hospital) using standard heat-shock protocol to obtain GFP-expressing *E. coli* to visualize bacteria movement within mucus as previously described(41). The GFP-expressing *E. coli* and *L. rhamnosus* from glycerol stocks were cultured in EZROM media (gift from Kim Lewis lab at Northeastern University) overnight (16 hours) at 37°C while shaking at 220 rpm. One day before bacteria inoculation, fluid within the channels of the human duodenal chips was replaced with N-[N-(3,5-Difluorophenacetyl-L-alanyl)]-(S)-phenylglycine t-butyl ester (DAPT, Sigma-Aldrich 565770) and prostaglandin E2 (PGE2, Tocris 2296) supplemented differentiation medium (DM-RN+DAPT+PGE2, Table 1) without antibiotics. On the day of bacteria inoculation, the human duodenal chips were stained with 10 μg/ml of WGA and 5 μg/ml of Hoechst 33342 for 30 minutes at 37°C. The *L. rhamnosus* was also stained with Hoechst 33342 (5 μg/ml) for 30 minutes at 37°C, followed by centrifuging at 12700 rpm at 4°C. Then, the stained *L. rhamnosus* pellet was resuspended with DPBS supplemented with 10 mM HEPES (Gibco™). Both the GFP-expressing *E. coli* and stained *L. Rhamnosus* were prepared in DPBS supplemented with 10 mM HEPES at a density of 5.7 × 10^6^ cells/ml and introduced into the lumen channel. Bacteria were co-cultured with the duodenal chips for 30 minutes at 37°C before imaging. A ZEISS LSM800 confocal microscope was utilized to collect a time-lapse video and z-stack fluorescence images of bacteria interactions with the mucus barrier.

### Statistical analysis

Differentiation medium formulation optimization was completed in three separate experiments, and the images shown in the results section were selected as representative images. ABT-450 amorphous nanoparticle tracking experiments were completed in triplicate, where at least 100 particle trajectories were analyzed in each experiment (on the mesofluidic gut model and native pig intestinal mucus model). Native pig intestinal mucus used in the particle tracking experiments was collected from three separate animals to account for biological variability. All quantitative data are presented as a mean value with standard error of the mean (SEM), and ANOVA test was used to determine significance with α=0.05.

## Result and discussion

### Development of mesofluidic gut chip enabling facile visualization of mucosal interface

A mesofluidic gut chip was designed and developed in which primary intestinal cells grow on a vertical hydrogel wall so that the mucosal interface can be directly visualized using a standard confocal microscope (Fig. 1). The mesofluidic gut chip consists of three channels, including a lumen channel representing the intestinal lumen, a side channel representing circulatory flow, and a stromal gel guide channel holding the vertical gel wall which separates the lumen and side channel (Fig. 1). Various natural and synthetic hydrogel materials, including gelatin methacryloyl (GelMA), type I collagen, and polyethylene glycol (PEG)-based hydrogels, were evaluated to test ability of these materials to: 1. form a vertical gel wall assisted by phase guides within the PDMS device, and 2. support primary organoid-derived human duodenal epithelial cell attachment and monolayer formation. We identified that the type 1 collagen gel wall could support attachment and culture of primary epithelium but was prone to deformation and subsequent leaking between channels, potentially due to forces generated by cultured cells or exerted during medium exchange (Supplementary Fig. 1A). A cross-linking procedure was thus adapted from previous reports of primary epithelial culture on collagen gel within a cell culture insert(35, 36). EDC-NHS cross-linking of one side of the type I collagen gel wall to increase mechanical integrity allowed it to still support formation of a primary human duodenal epithelial cell monolayer and also maintain gel wall integrity throughout the cell culture period (up to 10 days of culture). Within the mesofluidic gut chip, primary human duodenal epithelial cells formed a confluent monolayer on the collagen gel wall within 3 to 5 days after seeding in duodenal expansion media (EM) (Supplementary Fig. 1B). Attempts to visualize the mucus utilizing fluorescently conjugated WGA revealed a layer approximately 5 μm thick covering the duodenal epithelium after 5 days of differentiation in differentiation media (DM) (Table 1, Supplementary Fig. 1C and D).

**Fig 1.**
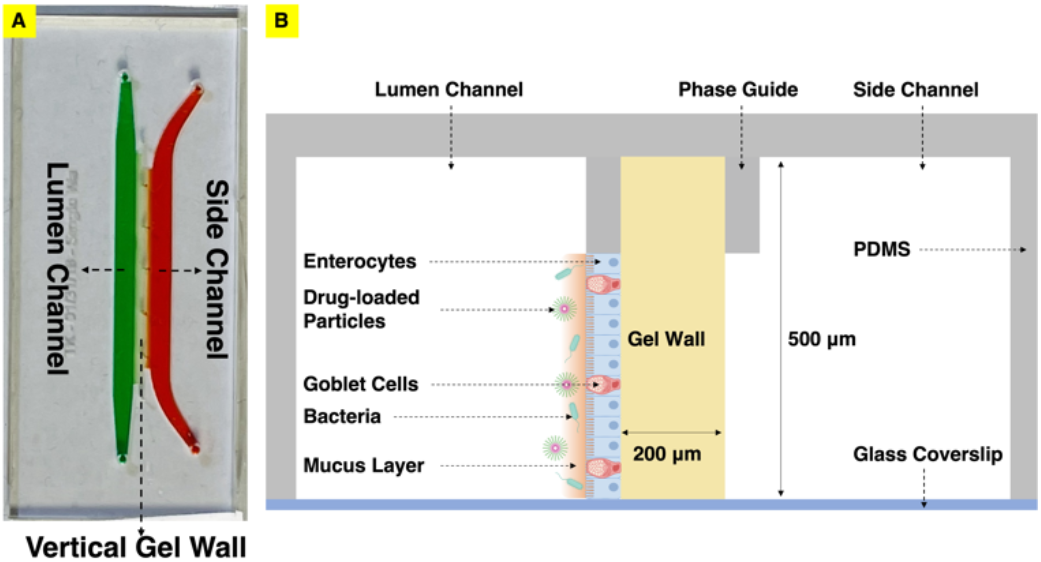
Mesofluidic gut model enabling visualization of the mucosal interface. (A) Top view, showing 3 channels including a lumen channel in green representing the intestinal lumen, a side channel in red representing circulatory flow, and a gel wall separating them. (B) Schematic cross-sectional view. The apical surface of primary human intestinal epithelial cells cultured on the gel wall faces the lumen. Luminal stimuli (bacteria, particulates, etc.) can be introduced into the lumen channel for studying their interactions with the mucosal barrier.

### Differentiation media optimization to improve robustness of human mesofluidic duodenal chip culture and increase mucus layer thickness

While primary human duodenal epithelial cells could form a confluent monolayer reproducibly on the cross-linked collagen gel wall, the duodenal monolayer in the majority of the chips delaminated after 3 days of differentiation in DM, leading to insufficient time for downstream experiments. In addition, a mucus layer close to physiological thickness (on the order of 100 µm)(8) was desired to enable utilization of the mesofluidic gut model to study bacteria and drug carrier transport within (e.g., via particle tracking) and penetration through an intact mucus layer. Hence, the differentiation media formulation was varied to increase the lifespan of duodenal monolayers and promote mucus production. In intestinal organoid and primary intestinal cultures, cells are generally maintained in a medium for some time containing essential factors present in the stem cell niche for promoting survival of intestinal stem cells, allowing for cell proliferation and expansion(18). This “expansion” period of culture typically lasts 2-5 days and is followed by removal of stem cell supporting factors from medium or addition of signaling small molecules in medium, allowing for differentiation of cells. This “differentiation” period of culture can last weeks(19, 29, 36, 42), depending on the robustness of the particular system, and is critical to differentiation of cells into specified epithelial cells, including mucus-secreting goblet cells required for quantitative analysis of mucus barrier properties. When organoids are used to seed cells on a culture surface (rather than being propagated as organoids within gels), the culture time is limited, as delamination of cells from a culture surface can occur after a certain period of differentiation, and this period of time is dependent on culture conditions including the substrate, medium composition, and medium volume. Wnt3a, R-spondin, and Noggin are critical factors for intestinal crypt cell proliferation, and Wnt3a, in particular, is an essential component among these three factors(43-45). It has recently been demonstrated that adding signaling molecules instead of removing WRN factors from medium results in robust differentiation of mature intestinal epithelial cell lineages in intestinal organoids(46). Hence, we hypothesized that providing R-spondin and Noggin in differentiation medium would enhance human duodenal monolayer viability over multiple days of culture while not sacrificing differentiation effectiveness(18, 47) as reflected in mucus production. Differentiation media supplemented with R-spondin and Noggin (DM-RN) was indeed determined to consistently maintain the integrity of human duodenal monolayers within gut chips throughout the 5-day differentiation period (Fig. 2); however, the lectin-labeled mucus layer upon these monolayers remained approximately 5 μm thick (Fig. 3).

**Fig 2.**
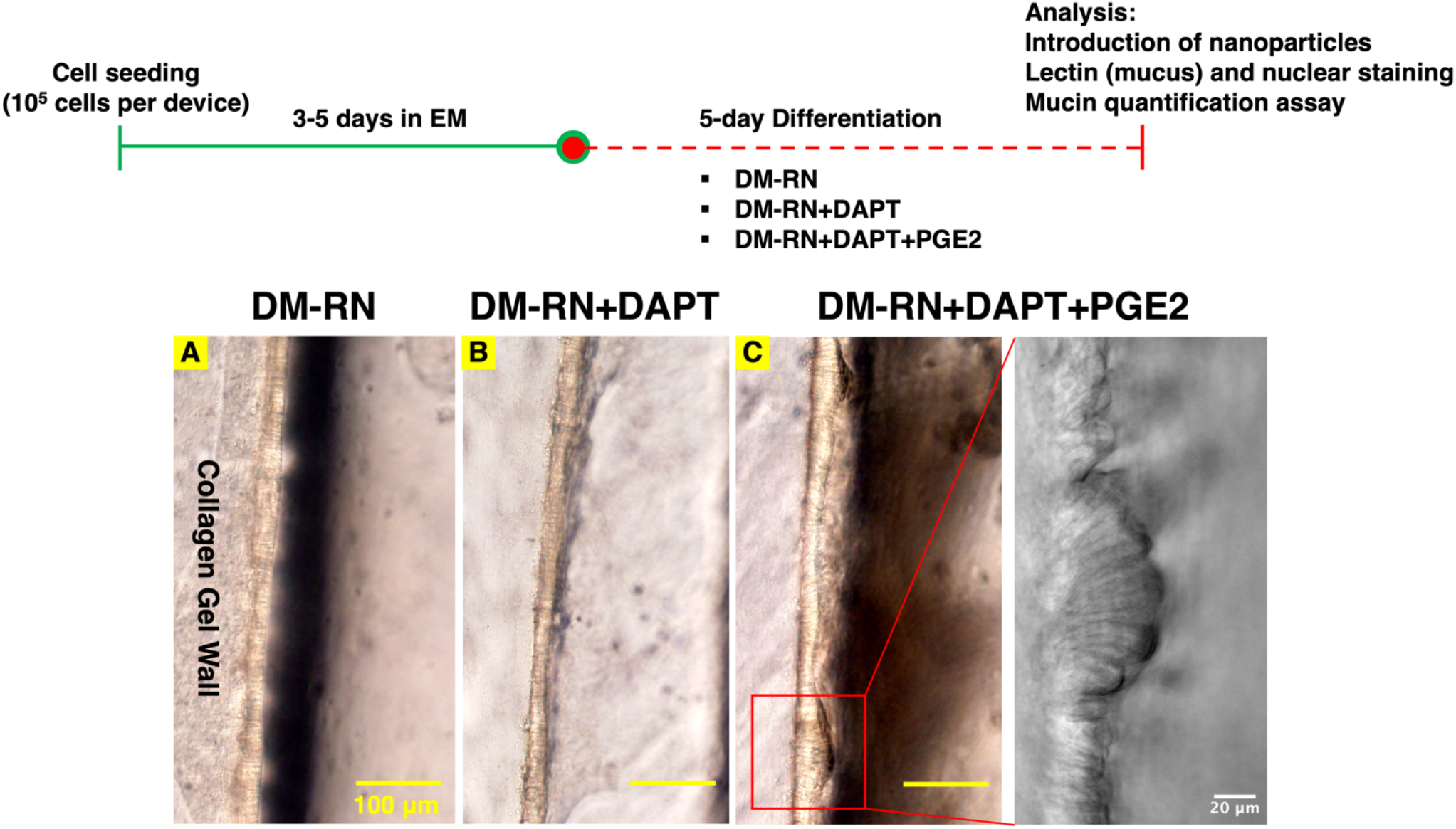
The human mesofluidic duodenal chips differentiated using modified differentiation media (DM-RN+DAPT+PGE2) demonstrate taller cell height with columnar shape of epithelial cells. Top: Human mesofluidic duodenal chip culture procedure and experimental timeline. Bottom: Brightfield microscopic images of primary human duodenal monolayers in the mesofluidic chips differentiated for 5 days using (A) differentiation media with R-spondin and Noggin (DM-RN), (B) DM-RN supplemented with 10 μM DAPT, and (C) DM-RN supplemented with 10 μM DAPT and 1.4 nM PGE2, with a higher magnification image showing the human duodenal epithelium morphology.

**Fig 3.**
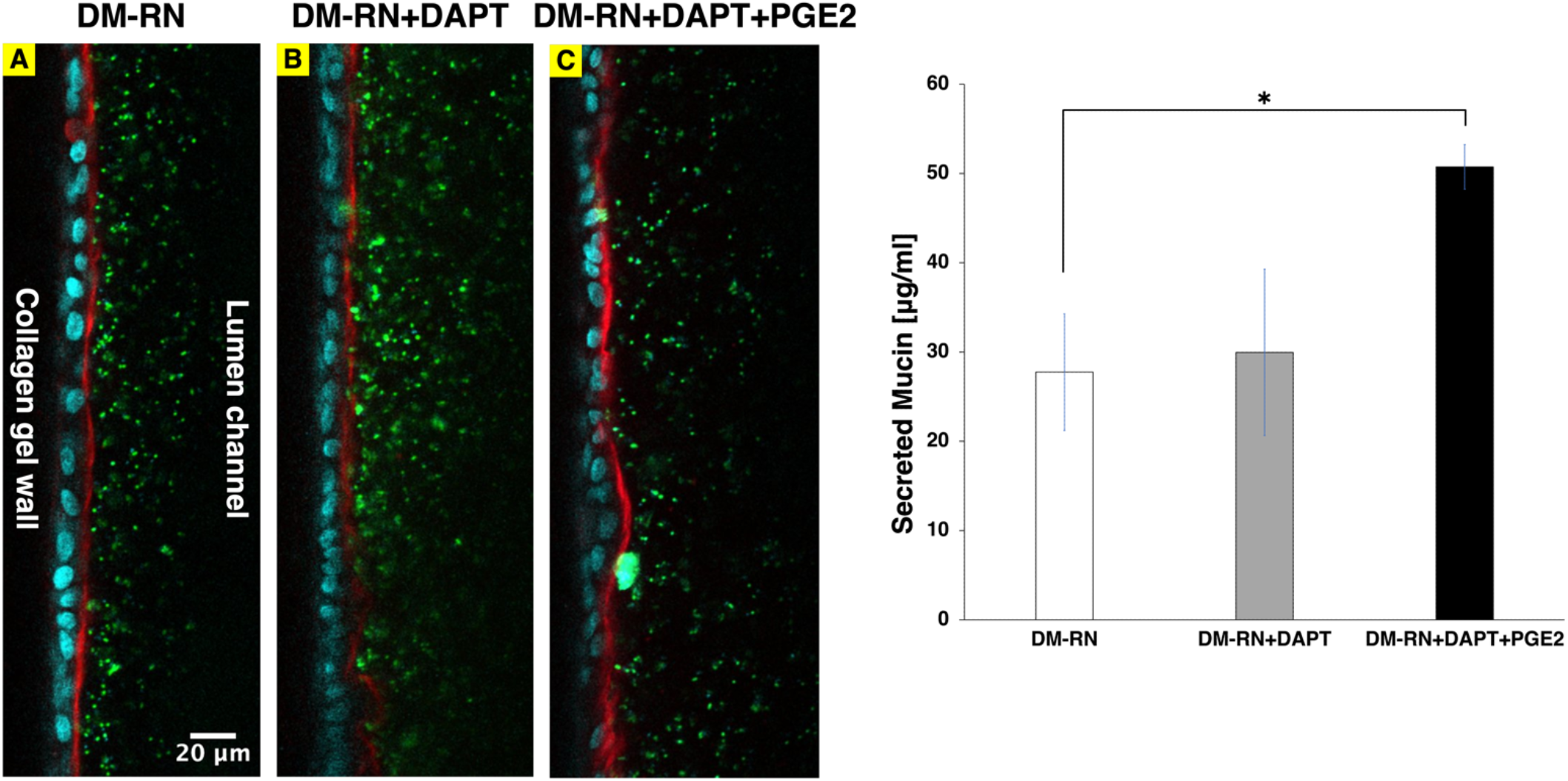
Left: Addition of DAPT and PGE2 during duodenal epithelium differentiation moderately increased mucus production. Primary human duodenal epithelial cells in the mesofluidic gut model were differentiated for 5 days using (A) differentiation media with R-spondin and Noggin (DM-RN) (B) DM-RN supplemented with 10 μM DAPT, and (C) DM-RN supplemented with 10 μM DAPT and 1.4 nM PGE2. Confocal fluorescence images are shown of the duodenal monolayers stained with fluorescently tagged lectin (WGA, red) for mucus and Hoechst 33342 (blue) for nuclei, with 200-nm fluorescent polystyrene microspheres (green) introduced to the lumen channel for visualizing their distribution relative to the edge of the mucus layer. Right: Alcian Blue mucin quantification assay showing moderate enhancement in secreted mucin in the duodenal chips differentiated using DM-RN supplemented with DAPT and PGE2 relative to DM-RN alone. * Indicates α < 0.05.

It has been shown that addition of DAPT in cell culture media could promote the differentiation of human intestinal organoids and increase the MUC2 mRNA expression in rectal and ileal organoids, indicating that DAPT might promote mucus production(18). Moreover, Sontheimer-Phelps et al. demonstrated that 6-day treatment with PGE2 on a human colon chip increased MUC2-expressing cells and proliferating cells, suggesting that PGE2 might also promote mucus production and aid in maintenance of duodenal monolayers in culture. Hence, the effects of DAPT and PGE2 on mucus production and duodenal monolayer integrity in the mesofluidic gut model were studied. Human mesofluidic duodenal chip cultures were seeded and cultured for 3-5 days (until confluent monolayer observation) in EM, followed by differentiation using three differentiation media formulations: DM-RN, DM-RN+DAPT, and DM-RN+DAPT+PGE2 (Table 1, Fig. 2). Duodenal monolayers differentiated using DM-RN or DM-RN+DAPT+PGE2 reproducibly maintained their integrity throughout the 5-day differentiation period (Fig. 2), while approximately 20% of the human duodenal chips differentiated using DM-RN+DAPT delaminated after 3 days of differentiation (Data not shown), indicating that maintenance of an intact barrier is enhanced by R-spondin, Noggin, and PGE2. Moreover, duodenal monolayers differentiated using DM-RN+DAPT+PGE2 exhibited taller columnar epithelial cells and a less flat, more variable structure compared to those differentiated with DM-RN or DM-RN+DAPT (Fig. 2). Confocal fluorescence imaging of cultures stained with fluorescently tagged lectin (WGA) showed that supplementation of DM-RN with DAPT and PGE2 moderately increased mucus layer thickness in the mesofluidic gut model from approximately 5 μm to 15 μm at the thickest part (Fig. 3). Fluorescent microbeads (200-μm carboxylate-modified polystyrene) introduced to the lumen channel filled the luminal space up to the lectin-stained layer, suggesting that the lectin was appropriately labeling the mucus layer (i.e., there was not a more diffuse layer that was not labeled with lectin but would still exclude particles). In addition, spent media collected from the duodenal chips differentiated with DM-RN+DAPT+PGE2 contained the highest mucin concentration, which was ∼2-fold greater than that measured in media from chips differentiated using DM-RN (Fig. 3), consistent with our microscopic observations (Fig. 3).

### Visualization of mucin turnover on chip

The ability to visualize the intact mucus layer on live cell culture within the mesofluidic gut model prompted attempts to capture dynamics of formation and potential turnover of the mucus layer. Imaging was thus performed approximately 16 hours after staining with lectin in an attempt to capture these processes. We predicted that mucus produced by the epithelium after staining would not be labeled with the lectin, while the outermost part of the mucus layer would be labeled by fluorescently tagged WGA. Remarkably, imaging 16 hours after staining revealed significant intracellular fluorescent signal, with the edge of the plasma membrane also clearly labeled (Supplementary Fig. 2), suggesting mucin turnover. A similar staining pattern was observed *in vivo* in rodents after labeling of mucus using *N*-azidoacetylgalactosamine (an analog of *N*-acetylgalactosamine), and was attributed to the early stages of mucin turnover(48).

### Air-liquid interface differentiation significantly increases the mucus layer thickness in the mesofluidic gut model

To generate a mucus layer with a more physiologically relevant thickness using our mesofluidic gut model, the epithelium was cultured at an air-liquid interface (ALI) during differentiation using DM-RN+DAPT+PGE2 in the side channel. ALI culture has been demonstrated to promote mucus production in airway epithelium cultures(49). Moreover, ALI culture was previously exhibited to enable production of a thick (>50 µm) mucus layer in human colonic epithelial cultures on cell culture inserts, and it was hypothesized that this was due to minimization of mucus dilution by apical media(50). In our mesofluidic duodenal chip cultures, maintaining apical ALI could also minimize mucus loss due to repeated medium changing. ALI differentiation might also minimize the dilution of signaling factors secreted apically, which may be critical to intestinal epithelial cell differentiation. ALI culture increased thickness of the mucus layer produced from approximately 5 µm to above 10 µm (Fig. 4). In ALI human colonic epithelial cultures on cell culture inserts, vasoactive intestinal peptide (VIP) incorporation into basolateral media led to a hydrated mucus layer of uniform thickness(50). VIP is an endogenous intestinal neuropeptide that can regulate intestinal barrier homeostasis(51), promote apical water secretion(50), and increase mucus secretion(52). To test if VIP could increase mucus layer thickness in our system, human mesofluidic duodenal chips were differentiated under either an ALI or submerged (i.e., with media in both channels) condition, with or without the addition of VIP. Inclusion of VIP increased the mucus layer produced in ALI cultures to approximately 50 μm after 5 days of differentiation (Fig. 4). The intact mucus layer served as a significant barrier to 200-nm fluorescent nanoparticles, evident by few of the nanoparticles penetrating the mucus layer after a 30-minute incubation. Measurement of mucin concentrations in spent media from the lumen compartment revealed that ALI+VIP increased secreted mucin levels by approximately 2.6-fold compared to submerged+VIP cultures (Fig. 4). While ALI without VIP resulted in a slight visible increase in mucus production, duodenal chips differentiated under the submerged condition with and without VIP had similar mucus production (Fig. 4), indicating an important synergistic effect of VIP and ALI on mucus production in our system. Human mesofluidic duodenal chips differentiated at ALI with DM-RN+DAPT+PGE2+VIP were subsequently utilized to study the transport of drug-rich amorphous nanoparticles (ANPs) and bacteria within the mucus layer, as described in the following sections.

**Fig 4.**
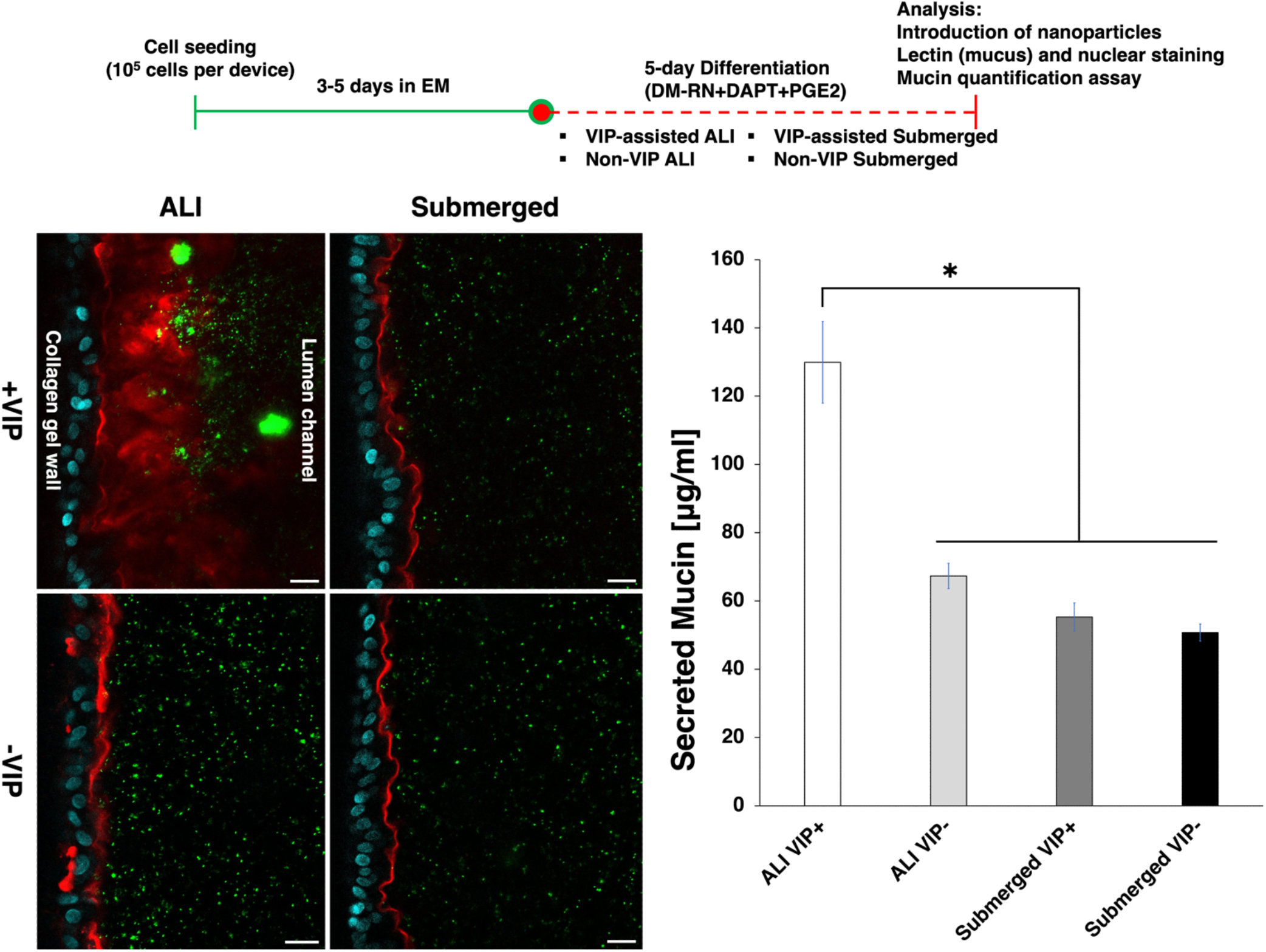
ALI differentiation with basolateral VIP exposure significantly increased mucus layer thickness. Top: Human mesofluidic duodenal chip culture procedure and experimental timeline. Bottom Left: The human mesofluidic duodenal chips were differentiated using 4 distinct methods: 1. + VIP at ALI, 2. +VIP submerged, 3. -VIP at ALI, 4. -VIP submerged. Human duodenal epithelium was then stained with WGA (red) for mucus and Hoechst 33342 (blue) for nuclei. 200-nm fluorescent polystyrene microspheres were introduced into the mesofluidic duodenal chips for visualizing their distribution within the lumen vs. at the edge of the mucus layer. Scale bar: 20 μm. Bottom Right: Alcian Blue mucin quantification assay showing significantly greater secreted mucin concentration in the duodenal chips differentiated using ALI+VIP compared to those differentiated using ALI without VIP and submerged conditions. * Indicates α < 0.05.

### Introduction of drug-rich amorphous nanoparticles in the mesofluidic gut model for studying their transport within an intact mucus layer

One motivation for developing the mesofluidic gut chip described here was to enable facile visualization of interactions between luminal contents, such as particulate drug carriers, and the mucosal interface. Amorphous nanoparticles (ANPs) are engineered drug rich structures which resemble *in situ* colloidal particles believed to be formed upon oral dosing of amorphous solid dispersion (ASD) formulations(39). ANPs have recently been reported in design of high drug loading amorphous formulations that overcome the high pill burden limitations of conventional ASDs(39). Analysis of the transport properties of ANPs and other drug-loaded particles within the intact mucus layer could offer novel insight into important factors to consider in drug delivery system development. Two formulations of drug-loaded ANPs prepared with Copovidone as the polymer and SDS as the surfactant: small ANPs (size: 125 nm, zeta potential: -43.8 ± 3.4 mV) and large ANPs (size: 230 nm, zeta potential: -47.6 ± 3.2 mV) were incubated within the mesofluidic gut model differentiated under the ALI+VIP condition with DAPT and PGE2 supplemented differentiation media (DM-RN+DAPT+PGE2+VIP in the side channel) or in *ex vivo* porcine intestinal mucus (Table 2) for 90 minutes. The multiple particle tracking (MPT) technique was then utilized to investigate transport of ANPs within native porcine intestinal mucus or the intact mucus layer of the mesofluidic gut model. Mean squared displacement (MSD) and effective diffusivity (D_eff_) were calculated from more than 100 particle trajectories in each experiment to quantify particle transport. The increase in particle size from 125 nm to 230 nm notably hindered particle diffusion within collected porcine intestinal mucus, indicated by reduced effective diffusivity from 0.051 to 0.032 μm^2^/s at a timescale of 3 seconds (Fig. 5). Similarly, larger particles diffused more slowly than smaller ones on the duodenal chip (0.047 vs. 0.07 μm^2^/s) (Fig. 5). Representative particle trajectories also reflect relatively unconstrained particle diffusion of small ANPs within the intact mucus layer in the mesofluidic gut model (Fig. 5). Size selectivity of the intact mucus layer in the duodenal chip and qualitative similarity with respect to measured diffusivity on chip and within native porcine intestinal mucus supports the mesofluidic gut model’s utility for studying the mucus barrier using the multiple particle tracking technique. Slightly higher values of diffusion coefficients of particles on the chip relative to those in native collected mucus could reflect the fact that mucus on the chip is bathed in fluid, interspecies differences in mucus barrier properties (human vs. porcine), or the impact of the porcine mucus collection process (e.g., scraping of some cellular material in addition to mucus, since cellular lipid and DNA content may impact mucus viscosity and barrier properties(3, 16, 53)). Differences in diffusion coefficients between collected mucus and mucus on chip could also be related to the fact that particle diffusion on the chip was visibly excluded from certain regions of WGA-stained mucus (Supplemental Video 1 and 2). Collection of mucus by scraping may result in mixing of different regions of the mucus layer such that observed barrier properties reflect properties of portions of mucus from which particles may be excluded *in vivo*.

**Table 2.**
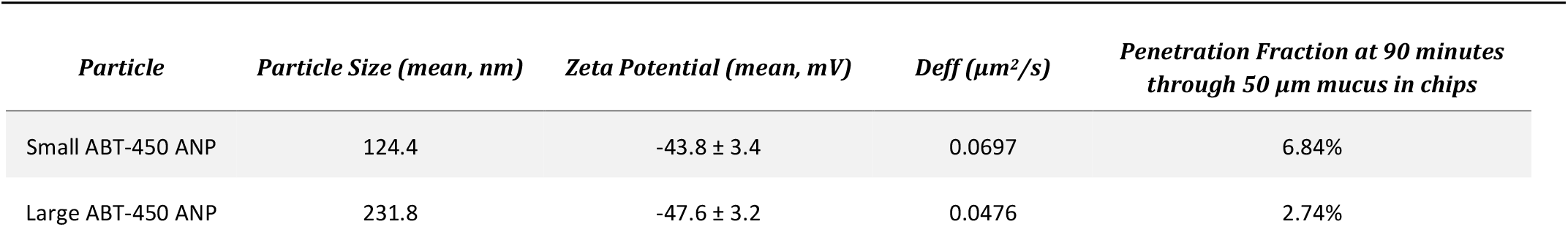
Prediction of Penetration Fraction of ABT-450 ANPs in Human Duodenal Chips.

**Fig 5.**
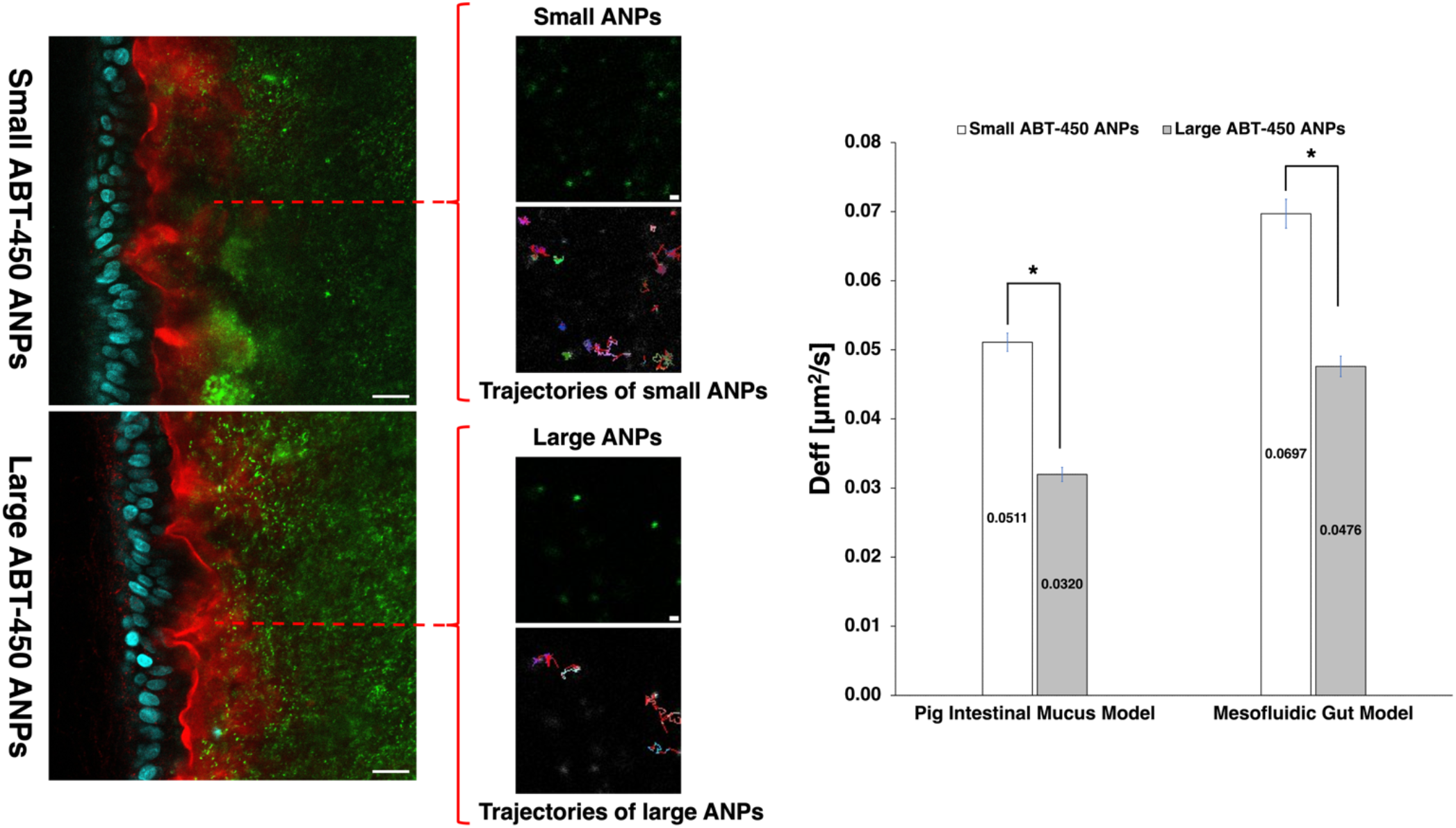
Multiple particle tracking technique was utilized to analyze diffusion of small and large ABT-450 ANPs in *ex vivo* collected porcine intestinal mucus and in the mesofluidic duodenal chips. Left: 20-second videos were collected at the regions of interest (within the WGA-stained intact mucus layer, pointed by dashed red lines) and particle trajectories were tracked using the Matlab algorithm. Representative particle trajectories of small and large ABT-450 ANPs are shown here. Long and short white scale bars: 20 and 1 μm, respectively. Right: Effective diffusivities of ABT-450 ANPs calculated using mean-squared displacement (MSD) from particle trajectories at τ = 3 sec indicated that larger particles have decreased effective diffusivity and particles diffuse more freely in the intact mucus layer relative to *ex vivo* porcine mucus. * Indicates α < 0.05.

Understanding the penetration rate of particles with different formulations through the intact mucus layer might provide insight into important parameters to consider in the design of drug delivery systems. To capture an advancing front of ANPs as they travel through the intact mucus layer in the mesofluidic gut model, fluorescence videomicroscopy was utilized to record time-lapse videos (30-second interval, 30-minute duration) at select time points (1.5 and 4 hours after particle introduction). The confocal fluorescent images collected at the beginning and end of the first recorded time interval: 1.5 hours and 2 hours after particle introduction, showed that the diffusing fronts of ANPs for both sizes of particles did not visibly advance toward the duodenal epithelium over the 30-minute time course (Fig. 6B, Supplementary Video 1). A significant fraction of ANPs had already penetrated to more than half the depth of the mucus layer at the beginning of the observation period (90 minutes after introducing particles). After the 90-minute incubation period, both small and large ANPs had diffused through approximately 3/4 depth of the intact mucus layer (Fig. 6B). A small fraction of small ANPs were observed to have reached the duodenal epithelium’s apical surface, a phenomenon not observed with large ANPs (Fig. 6B, white circles). There was a visible increase in the concentration of large ANPs within the mucus layer over the 30-minute observation period, especially within pockets of the mucus layer with low levels of lectin staining (Fig. 6B, yellow arrows), while there was no significant visible change in particle distribution for small ANPs over this time period (Fig. 6B). Similarly, the fluorescence time-lapse videos at 4 hr after particle introduction did not capture notable movement of the diffusing fronts of ANPs for either particle size (Supplementary Video 1). The penetration fraction of ABT-450 ANPs through a mucus layer of 50 μm thickness after 90 minutes can be predicted using the measured particle effective diffusivity in conjunction with a numerical integration of Fick’s second law. It was estimated that approximately 7% and 3% of small and large ABT-450 ANPs, respectively, would penetrate through the mucus layer of 50 μm thickness in 90 minutes (Table 2), which agrees reasonably with visible observations. For future study, efforts will include identifying the optimal incubation time that will allow visualization of the particle penetration process starting from the mucus layer edge to be captured.

**Fig 6.**
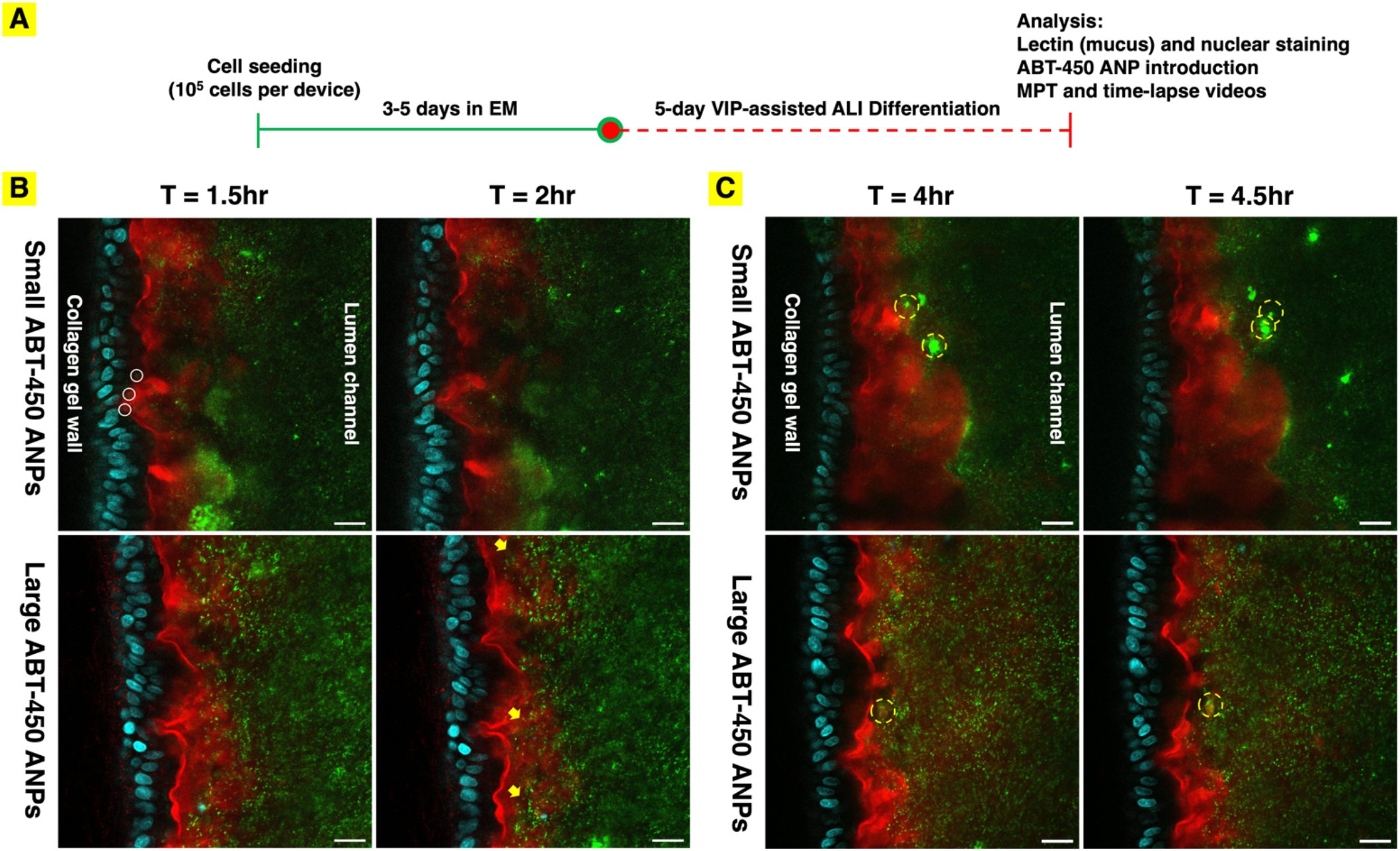
(A) Human mesofluidic duodenal chip culture procedure and experimental timeline (B) Distributions of ABT-450 ANPs within the mucus layer in human duodenal chips differentiated under the ALI+VIP condition 1.5 hrs after introduction to the lumen channel. Confocal fluorescence images of first and last frames of the time-lapse videos (30-second interval, 30-minutes duration) show changes in distribution and location of ABT-450 ANPs (green) of two different sizes (“small”=125 nm and “large”=230 nm) within the WGA-stained (red) mucus layer over the 30-minute of period. White circles show small ABT-450 ANPs reaching to the epithelium after 1.5 hour. Yellow arrows show increased concentration of large ABT-450 ANPs in the less WGA-stained regions over the 30-minute observation period. (C) Confocal fluorescence images collected 4 hrs after introduction to the lumen channel of ABT-450 ANPs (green) reveals clumping and shedding of some ANPs away from the epithelial surface over time. Dashed yellow circles highlight the same clustered ANPs shedding away from the mucus layer over 30-minute observation period. Epithelial nuclei stained with Hoechest33342: blue. Scale bar: 20 μm.

Interestingly, while net advancement of the particle diffusion front was not obvious at the chosen time intervals, net bulk motion of some particles away from the epithelial surface was. Observing the WGA-stained region in isolation over the time course of the time-lapse videos enabled visualization of the shedding of portions of the mucus layer into the lumen channel (Supplementary Video 2). This is of significant interest, as formation and shedding of mucus is known to occur *in vivo*, potentially limiting penetration of particulate materials and even molecular species that interact with mucus components to the underlying tissue. Indeed, the clumping and shedding of particles away from the epithelial surface, likely with mucus components, could be observed, especially 4 hours after introduction of ABT-450 ANPs (Supplemental Video 1 and 2, Fig. 6C).

### Visualization of E. coli and L. rhamnosus interaction with the mucosal interface

To validate the utility of the mesofluidic gut model for visualizing and studying interactions of bacteria with the mucus barrier and apical epithelial interface, GFP-expressing *E. coli* and Hoechst 33342 labeled *L. rhamnosus* were inoculated into the duodenal chips to establish short-term co-cultures with human duodenal epithelium. After 30 minutes of co-culture, bacterial movement in the lumen channel was recorded via time-lapse imaging (2-minute interval, 1-hour duration). Bacteria remained largely mobile over the period of observation, exhibiting transient interactions with mucus, and tended to migrate in clusters(54) (Supplementary Video 3, Supplementary Fig. 3). Confocal fluorescence images captured at selected time points demonstrated that bacteria moved in close proximity to the apical membrane of the duodenal epithelium and were largely excluded from regions of intense WGA staining over the imaging period (Supplementary Fig. 3). The co-culture period of human cells and bacteria can be extended in future studies to capture potential bacteria penetration through the epithelium, as it may be of interest in understanding how this process relates to lumen contents and host cell response. Moreover, with the co-culture of primary human intestinal cells and commensal microbial consortia over extended culture periods, the mesofluidic intestinal model can serve as a useful tool for studying gut-microbiome homeostasis.

## Conclusion

The developed human mesofluidic duodenal chip offers a novel platform for visualizing the mucosal interface and studying interactions between the intestinal mucus layer and luminal contents. Optimizing differentiation media and incorporating air-liquid interface culture enabled the formation of a physiologically relevant (i.e., 10s of microns thick) mucus layer on a live epithelial culture whose integrity could be maintained over the cell culture period. The chip offers the ability to visualize the spatial location of particles and bacteria within the mucus layer on a live culture such that quantitative analysis of their motion (e.g., using particle tracking) and their spatial relation to the epithelial surface could be analyzed, providing valuable insight into the ability of drug delivery systems to penetrate through mucus and host-microbe interactions. Further, interesting phenomena relevant to both drug delivery and mucosal biology were revealed using time-lapse imaging on live cultures, including apparent clumping of particles and clustering of bacteria within mucus, mucus shedding from the epithelial surface, and uptake within cells of mucus components. Future studies can leverage this platform to further explore the dynamics of mucosal barrier function and its implications for drug transport and host-microbe interactions.

## Acknowledgements

Research reported in this publication was supported by National Institute of Biomedical Imaging and Bioengineering (NIBIB) and National Institute of General Medical Sciences (NIGMS) of the National Institutes of Health under award numbers R01EB021908 and R01GM098117, respectively. The authors thank Joshua Luchan and Cuiying Zhang for assistance in developing and maintaining mesofluidic human duodenal chip cultures; Abigail Koppes and Guohao Dai for training and use of the plasma cleaner; Danielle Large and Debra Auguste for training and use of the zeta potential analyzer; Mengqi Ren and Ke Zhang for training and use of the rotary evaporator. The authors thank Guoxin Rong, Core Facility Director of Institute for Chemical Imaging of Living System (CILS) at Northeastern University, for training and assistance in troubleshooting imaging procedures.

