## Supplemental Fig. 1,2, and 3 for "Human Mesofluidic Intestinal Model for Studying Transport of Drug Carriers and Bacteria Through a Live Mucosal Barrier"

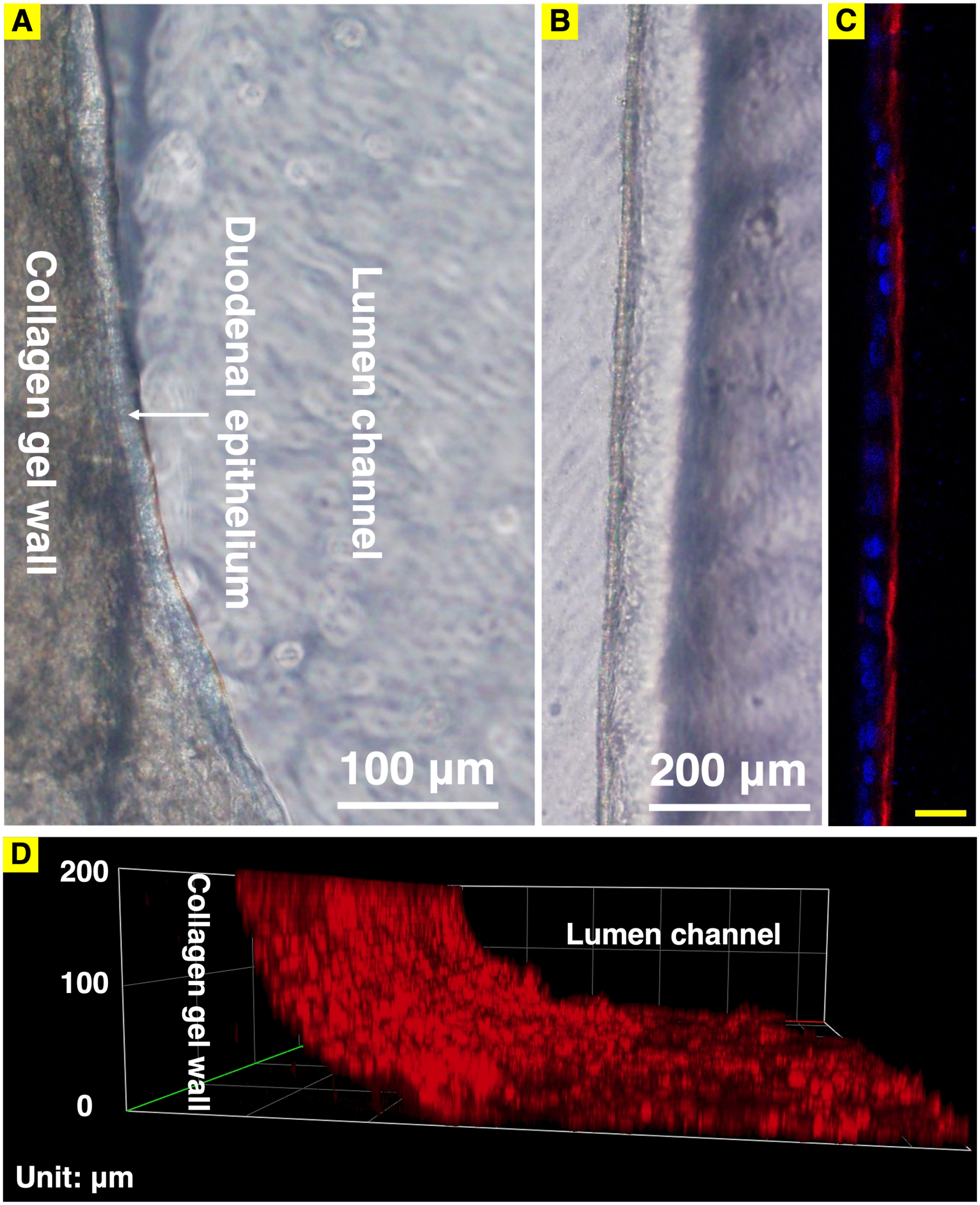


**Supplementary Fig. 1** Brightfield microscopic images showing (A) deformation of the non-crosslinked type I collagen gel wall and (B) maintenance of crosslinked type 1 collagen gel wall integrity after 4 days of culture in EM and formation of a confluent primary human duodenal epithelial monolayer. EDC-NHS crosslinking procedure increased the mechanical integrity of the collagen gel wall. (C) Fluorescence confocal image showing a thin mucus layer labeled by WGA (red) covering the human duodenal epithelium on the collagen gel wall differentiated for 5 days using DM.

Cell nuclei were stained by Hoechst 33342. (D) 3-D reconstructed fluorescence image of (C) showing a thin mucus layer covering the collagen gel wall to the height of 200 µm. Yellow scale bar: 50 µm.

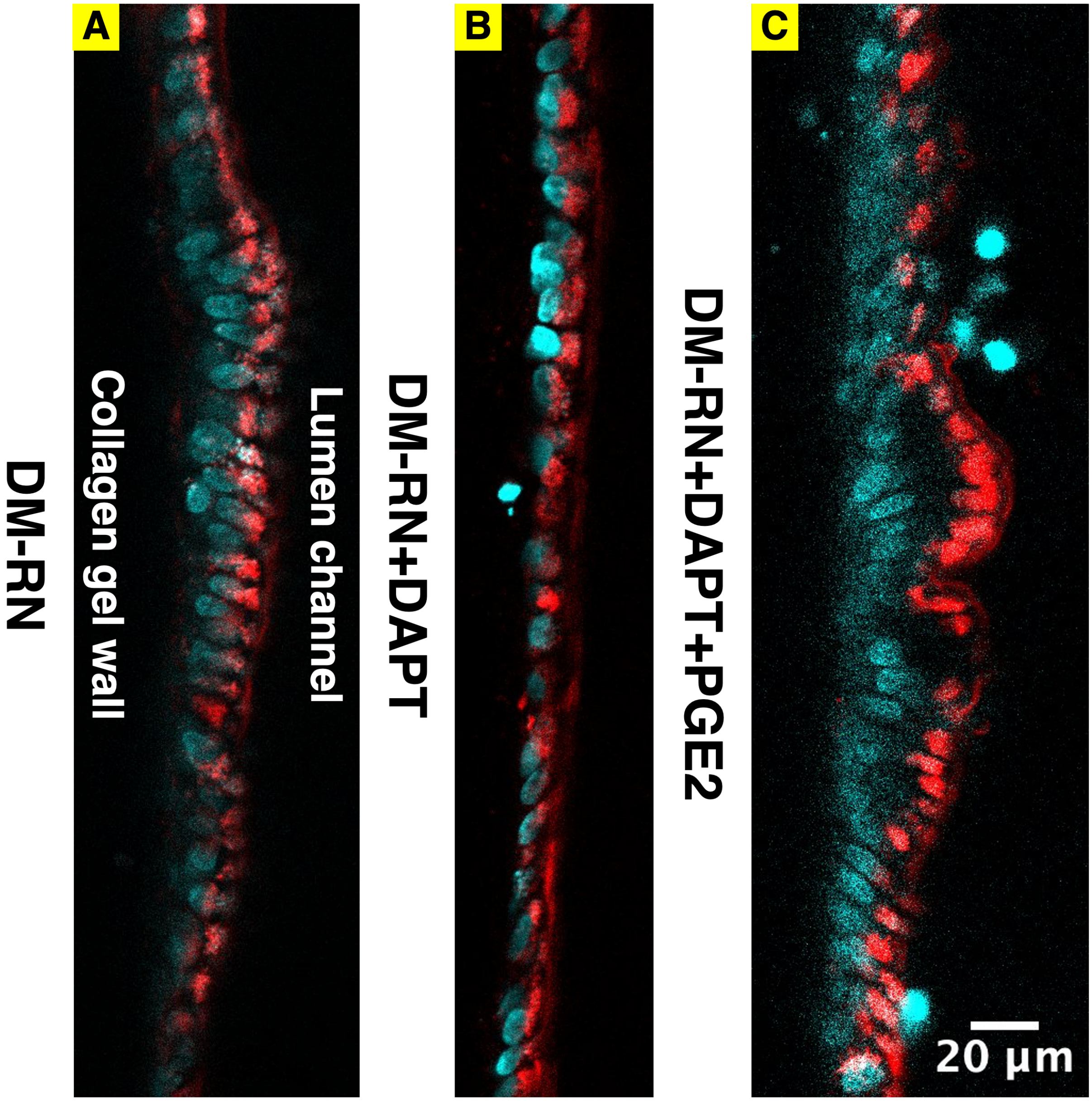


**Supplementary Fig. 2** Confocal fluorescence images of mesofluidic duodenal chips stained with WGA for mucus and Hoechst 33342 for nuclei one day (approximately 16 hours) before imaging. Intracellular WGA staining suggested uptake of lectin-stained material and potential mucus turnover by the duodenal epithelial cells after overnight incubation.

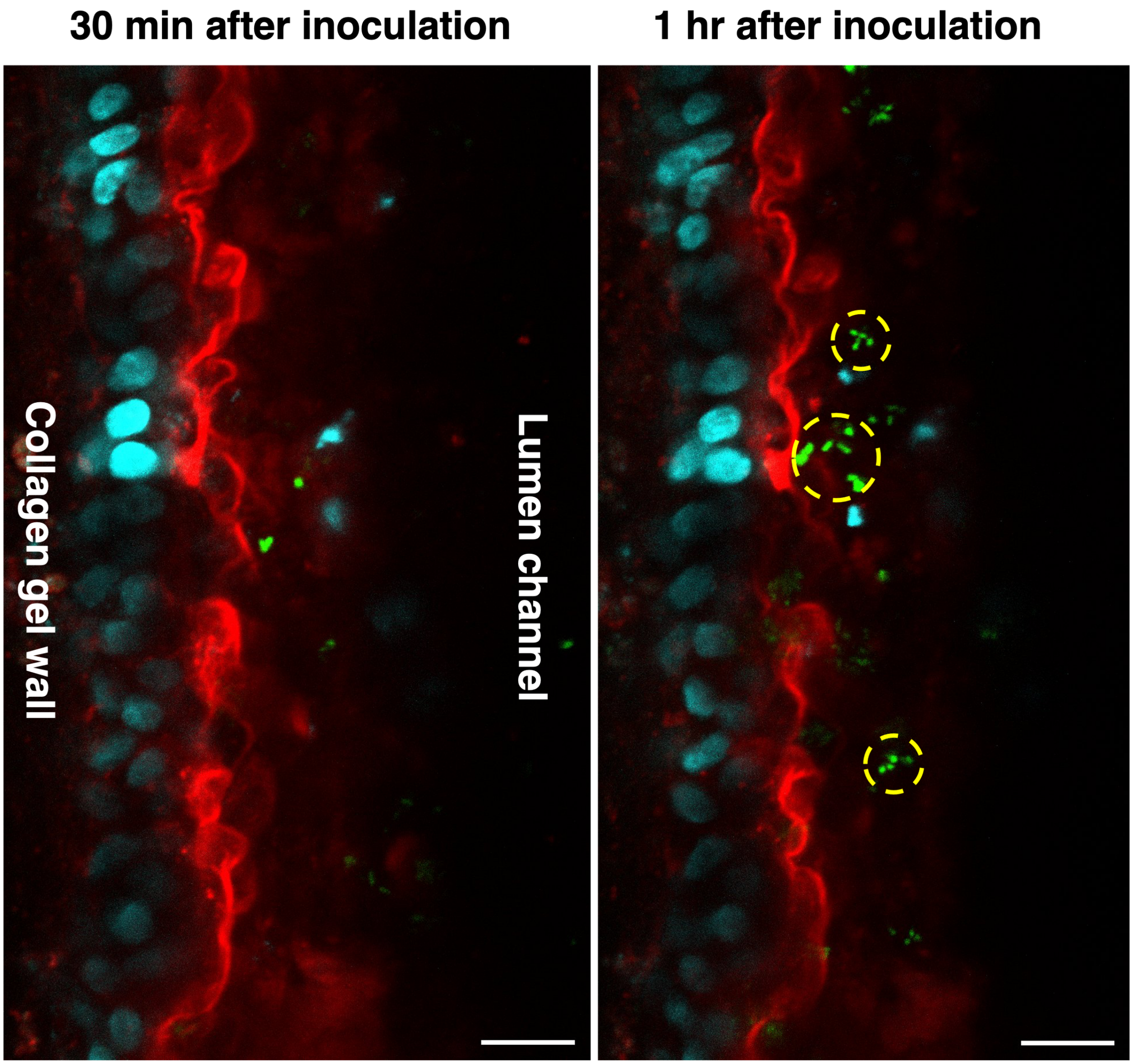


**Supplementary Fig. 3** Short-term co-cultures of the mesofluidic duodenal chips with bacteria. The human duodenal chips were stained with WGA (red) for mucus and co-cultured with GFP-expressing *E. coli* (green) and Hoechst 33342 labeled *L. rhamnosus* (blue). Fluorescence confocal images showing bacteria moving in clusters in close proximity to the duodenal epithelium over the 30 minute imaging period. *E. coli* moving in clusters are highlighted by yellow dashed circles. Scale bar: 20 µm.
